# Long-term dynamics of aberrant neuronal activity in Alzheimer’s disease

**DOI:** 10.1101/801902

**Authors:** V. Korzhova, P. Marinković, P. M. Goltstein, J. Herms, S. Liebscher

**Author notes:** **Highlights:** - “hyperactivity” is present in awake APPPS1 mice - activity levels of individuals cells in APPPS1 mice are largely stable, with small fluctuations comparable to those seen in WT mice - hyperactivity emerges largely from slow activity gain of intermediately active neurons - activity fluctuations are independent from plaque proximity, but aberrant activity levels are more likely to sustain in the plaque vicinity.

## Abstract

Alzheimer’s disease (AD) is associated with aberrant neuronal activity levels. How those activity alterations emerge and how stable they are over time *in vivo*, however, remains elusive to date. To address these questions we chronically recorded the activity from identified neurons in cortex of awake APPPS1 transgenic mice and their non-transgenic littermates over the course of 4 weeks by means of calcium imaging. Surprisingly, aberrant neuronal activity was very stable over time. Moreover, we identified a slow progressive gain of activity of former intermediately active neurons as the main source of new highly active neurons. Interestingly, fluctuations in neuronal activity were independent from amyloid plaque proximity, but aberrant activity levels were more likely to persist close to plaques. These results support the notion that neuronal network pathology observed in AD patients is the consequence of stable single cell aberrant neuronal activity, a finding of potential therapeutic relevance.

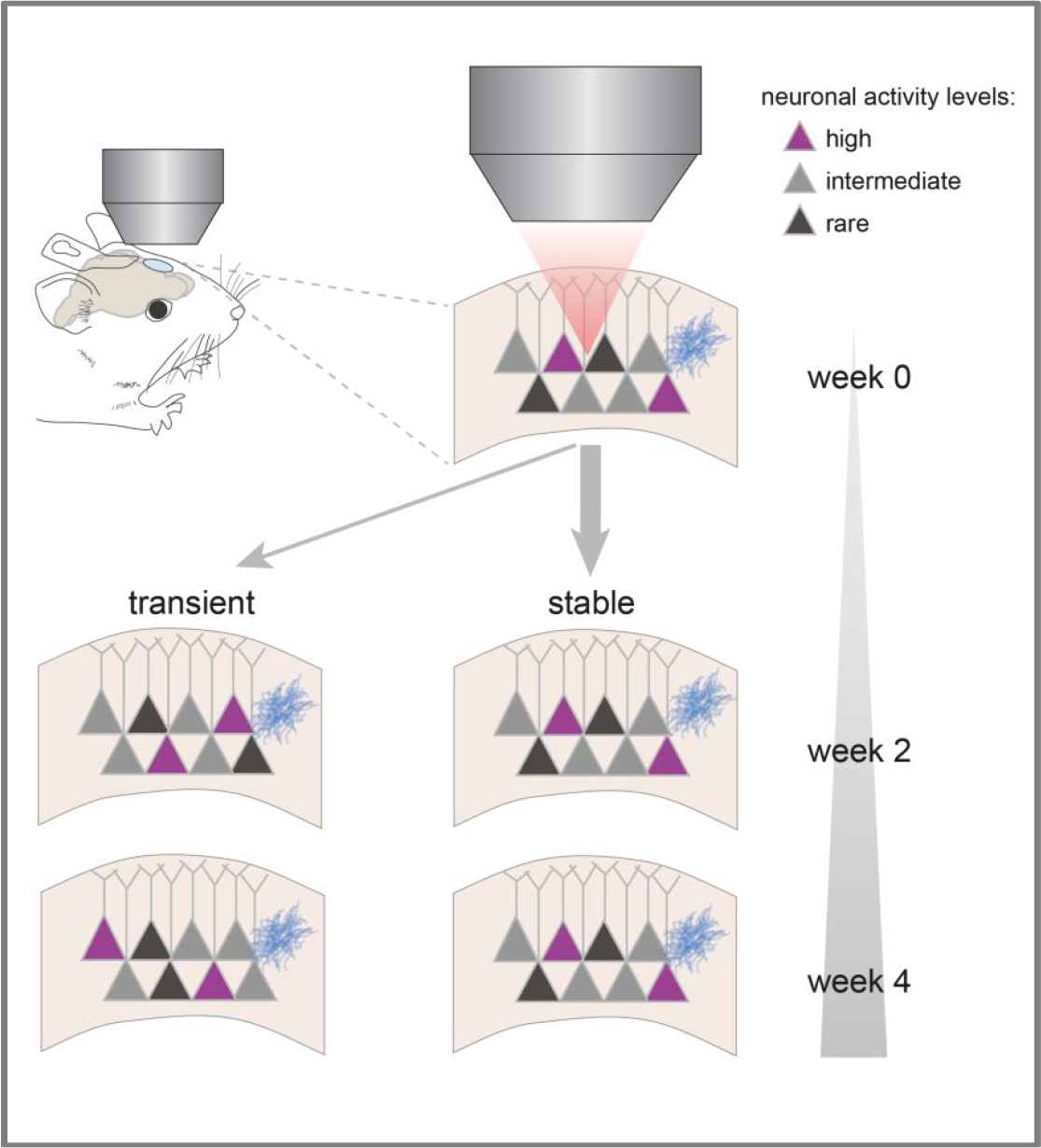

## Introduction

Alzheimer’s disease, the most common form of dementia, is histopathologically characterized by the accumulation of diverse assemblies of the amyloid–beta peptide (Aβ) within the CNS, which are accompanied by aberrant high and low neuronal activity levels, termed hyper- and hypoactivity ^1, 2^. Whether these aberrant activity levels are spatially confined to proteinaceous deposits, mainly consisting of Aβ, so-called amyloid plaques, remains a matter of debate ^1, 3, 4^. Importantly, aberrant neuronal activity has been shown to be one of the very early events of a pathogenic cascade, to worsen over time and thus to determine the level of cognitive impairment in affected individuals ^2, 5, 6^.

The mechanisms underlying the development of aberrant activity as well as their long-term dynamics *in vivo*, however, remain elusive thus far. Hyper- and hypoactivity of individual neurons could either be a transient phenomenon or alternatively represent a stable feature present over weeks or months. Both scenarios are conceivable given the different mechanisms suggested to underlie the development of these activity changes. Oligomeric Aβ has been suggested to directly or indirectly affect synaptic function. Aβ has been shown to bind to several surface molecules of synapses ^7, 8^, particularly to postsynaptic elements of excitatory synapses ^9^. Moreover, Aβ causes synaptic instability, a ‘shrinkage of spines’ and synaptic loss, which are accompanied by a reduction of LTP and increased LTD both *in vitro* and *in vivo* ^10–12^. Counterintuitively, Aβ can trigger an ‘aberrant’ increase in neuronal activity both acutely as well as under chronic conditions, such as in transgenic mouse models, which has been suggested to result from the activation of NMDA and AMPA receptors ^1, 13–16^ through increased glutamate release probability ^17^ or blockage of glutamate uptake ^18–20^. If direct stimulation of glutamate receptors was the main driver of aberrantly increased neuronal activity, one would expect at least initially a subsequent desensitization and/or downregulation of these receptors and consequently a normalization of the altered neuronal activity as a homeostatic response ^21^. Altered neuronal activity patterns in this case could occur rapidly and would be transient and dynamic in nature.

On the other hand, recent evidence suggests that different cell-types are affected differentially in the disease. As such, dysfunction of certain types of interneurons, e.g. parvalbumin -, somatostatin - or vasoactive intestinal peptide expressing interneurons, has been reported ^3, 22, 23^. These cell-type specific impairments are likely to cause a deficit of the otherwise tightly regulated balance between excitation and inhibition (E/I), leading to more global, circuit-level defects in AD. E/I imbalance could thus also underlie the observed changes in activity levels of single cells. This network remodelling, however, is likely to cause a more stable, slowly progressing change in neuronal activity. Insight into these single cell features are not only informative regarding the underlying pathomechanism, but, moreover, have potential therapeutic implications. A first critical step towards better understanding of the pathogenesis of activity changes in AD is to follow single neurons *in vivo* over extended periods of time.

## Results

To address this question, we longitudinally monitored the activity of individual neurons in layer 2/3 of the frontal cortex by means of calcium imaging using the genetically encoded calcium indicator GCaMP6s ^24, 25^ in awake Amyloid Precursor Protein – Presenilin 1 (APPPS1) ^26^ transgenic mice. We measured neural activity in three consecutive imaging sessions spaced by 2 weeks (Fig. 1a-c). We started our recordings at the age of 4.5 months, which in this mouse model represents the early phase of Aβ plaque deposition ^26, 27^. To characterize the cell identity of the imaged neurons, we performed post-hoc immunostainings and found that the vast majority (96.4 %) of GCaMP6s expressing cells was in fact excitatory (Suppl. Fig.1). We first compared the average neuronal activity of layer 2/3 neurons in frontal cortex of APPPS1 mice to their WT control littermates and found both the frequency of transients (Fig. 1d) as well as the area under the curve of the calcium signal, measured as an integral of the ΔF/F trace over time (Fig. 1e), to be significantly increased in APPPS1 mice throughout the experimental period.

**Fig. 1.**
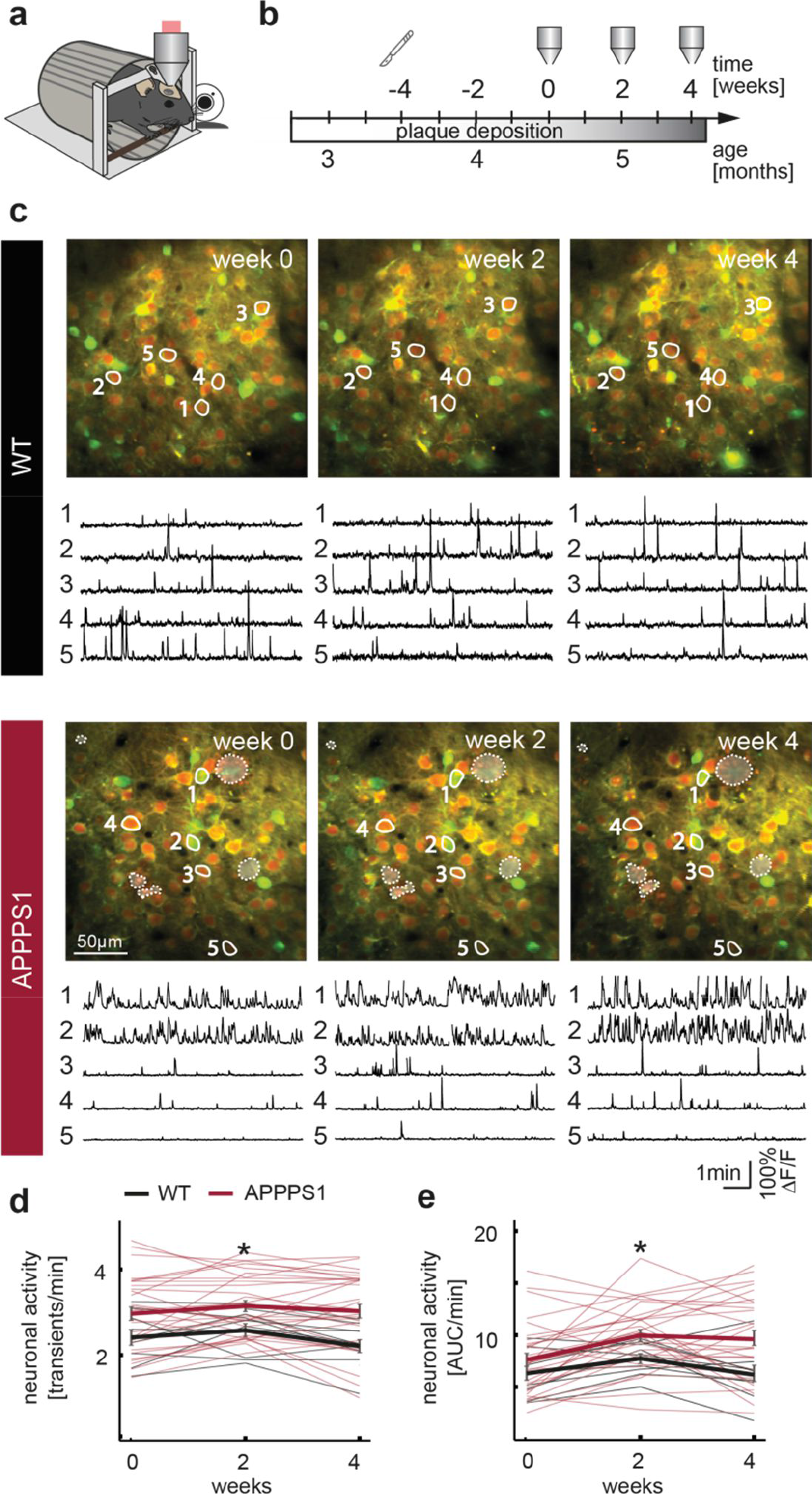
Chronic *in vivo* imaging of individual neurons in awake mice. **(a)** Schematic of the *in vivo* imaging setup. Mice were head-fixed and positioned in a restrainer, while neuronal activity was recorded through a cranial window above the frontal cortex. Whisking was recorded by a web camera. **(b)** Experimental timeline. Cranial window implantation (scalpel icon) was conducted four weeks prior to the first imaging session. Imaging (objective icon) was performed at two week intervals. Plaque deposition in the APPPS1 mouse line starts at ~8 weeks of age and plaque growth takes place throughout the life time of these mice (indicated by the gray gradient). **(c)** Representative average projections of a recorded field of views (FOV) in a WT (upper panel) and an APPPS1 (lower panel) mouse (mRuby2 expression shown in red, GCaMP6s in green). Example calcium traces of neurons labelled within the projections are shown below. Dashed lines denote the location of amyloid plaques. **(d)** Average neuronal activity of neurons in layer 2/3 in the frontal cortex of WT and APPPS1 mice, measured as transients per minute. Thin lines represent the averages of individual FOVs (same set of neurons imaged over three consecutive time points), thick lines represent the mean +/− SEM for each time point (effect of group: *F*_1,72_ = 6.07, *p* = 0.02; effect of time: *F*_2,72_ = 2.23, *p* = 0.12; group-by-time interaction effect: *F*_2,72_ = 0.91, *p* = 0.41, two way repeated measures ANOVA). **(e)** Neuronal activity measured as area under the curve (AUC) per minute across all time points (thick lines represent the mean +/− SEM, effect of group: *F*_1,72_ = 4.58, *p* = 0.04; effect of time: *F*_2,72_ = 9.72, *p* = 0.0002; group-by-time interaction effect: *F*_2,72_ = 1.67, *p* = 0.2, two way repeated measures ANOVA, WT n = 9, APPPS1 n = 29 experiments) is increased in APPPS1 mice. * *p* < 0.05

### Activity change over time

How do activity levels of individual neurons change over the time course of 4 weeks? To address this question we compared the distribution of the changes in activity of individual neurons between time points (Fig. 2a,b). Interestingly, the distribution of those activity changes did not differ between WT and APPPS1 mice between week 0 and 2, and was only slightly altered from week 0 to 4 (Fig. 2b). Moreover, we computed the similarity of activity levels of individual neurons within a given field of view across time points (Suppl. Fig.2) and did not observe a difference between WT and APPPS1 mice (Fig. 2c). To mimic a rapid and transient change in activity levels, we shuffled the data set by randomly changing the order of neurons. The similarity indexes of the shuffled data were close to zero and thus significantly lower than the similarity observed within the actual data set (Fig. 2c). Together these findings indicate that overall activity levels remain largely constant over time with only minor fluctuations in activity (+/− 2 transients/min for > 75% of all neurons). Thus aberrant activity is a stable single cell feature in AD.

**Fig. 2.**
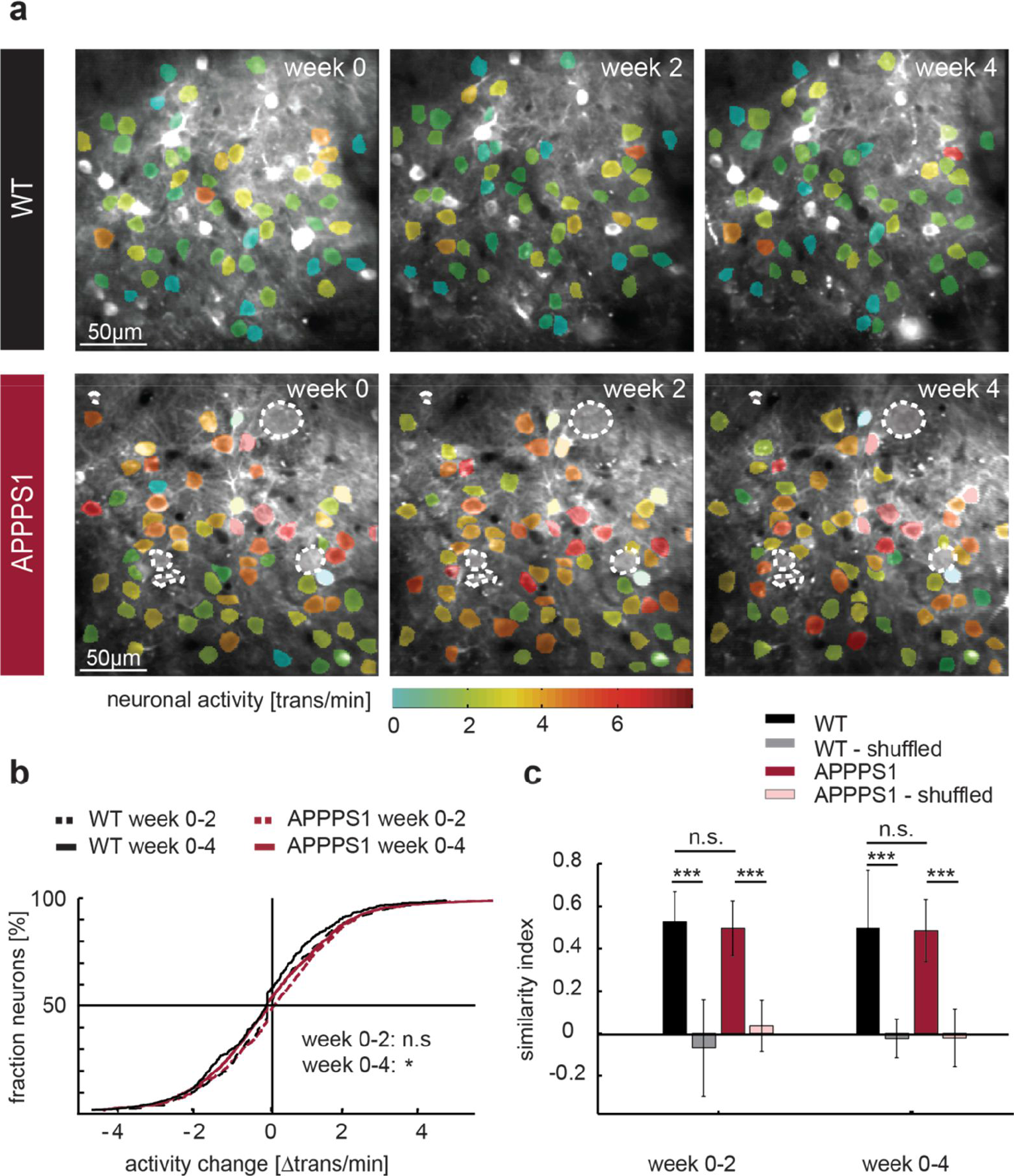
Neuronal activity levels are largely stable over time. **(a)** Representative mean projections of all 3 consecutive imaging time points from a WT (upper panel) and APPPS1 (lower panel) mouse, superimposed by the ROI (region of interest) selection mask, in which color codes for neuronal activity (transients per minute) - of note, filled cells were not included in the analysis. Dashed lines denote location of amyloid plaques. **(b)** The change in neuronal activity between week 0 – 2 and week 0 – 4 is shown (week 0 – 2: *p* = 0.08, week 0 – 4: *p* = 0.03, KS test, WT n = 562 neurons, APPPS1 n = 1691 neurons). **(c)** The similarity of neuronal activity within a given field of view is compared (week 0-2: *F*_3,74_ = 125.4, *p* < 10^−4^, one way ANOVA, WT vs WT shuffled *p* < 10^−4^, WT vs APPPS1 *p* = 0.9, APPPS1 vs APPPS1 shuffled *p* < 10^−4^, Bonferroni’s post-hoc test; week 0-4: *F*_3,74_ = 59.93, *p* < 10^−4^, one way ANOVA, WT vs WT shuffled *p* < 10^−4^, WT vs APPPS1 *p* = 0.9, APPPS1 vs APPPS1 shuffled *p* < 10^−4^, Bonferroni’s post-hoc test). Data are mean +/− SEM. * *p* < 0.05, *** *p* < 0.001

### Emergence and fate of highly active cells

We next asked whether the change in activity hinges on the initial activity level. To this end we classified neurons based on their activity levels into highly (>4 transients/min), intermediately (0.25-4 transients/min) and rarely (<0.25 transients/min) active neurons. In agreement with previous studies, conducted in anesthetized mice ^1,4^, we found that also awake APPPS1 mice had a significantly larger fraction of highly active neurons, while intermediately active cells were less abundant in APPPS1 mice (Fig. 3a,b). The fraction of rarely active cells, however, was not significantly affected in APPPS1 mice (Fig. 3a,b). Overall, the fractions of the different activity categories remained largely constant in both genotypes throughout the 4 week imaging period. A category-specific investigation, however, did reveal differences. As such highly active neurons in APPPS1 mice underwent a lower reduction in activity than their counterparts in WT mice (Fig. 3c). Intermediately active neurons in APPPS1 mice, on the other hand, displayed a small increase in activity over time compared to intermediately active neurons in WT mice (Fig. 3d).

**Fig. 3.**
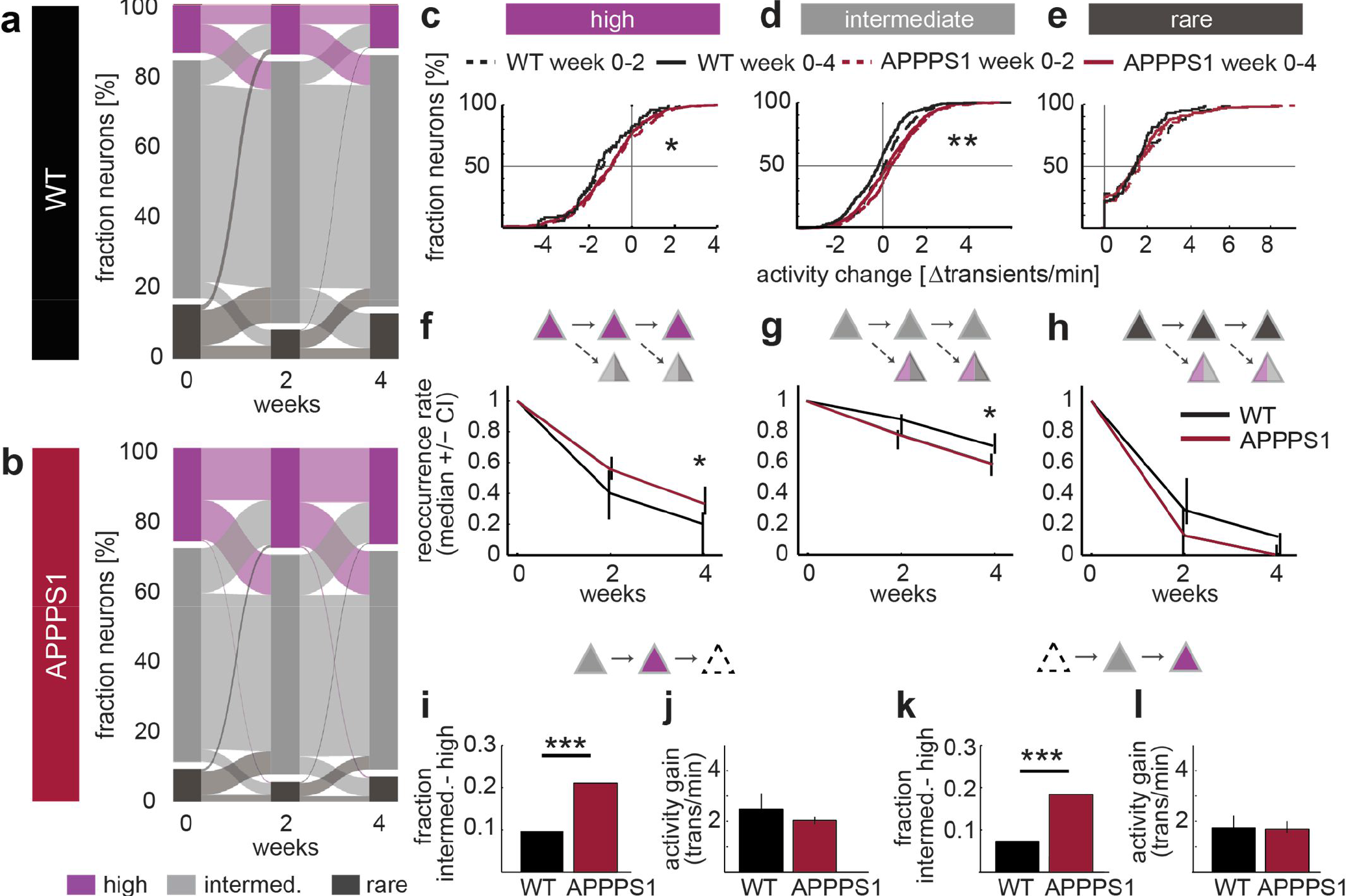
Category-specific changes in neuronal activity. **(a,b)** Alluvial plots depicting the fractional change of highly active (>4 transients/min), intermediately active (0.25 −4 transients/min) and rarely active (<0.25 transients/min) neurons in WT (A) and APPPS1 (B) mice over time (all neurons pooled in alluvial plots, fraction highly active neurons: week 0 – 4 in WT: 15.9 +/− 2.7%, 13.4 +/− 2.8 %, 10.6 +/− 2.5% and in APPPS1: 27.2 +/− 3.2 %, 27.7 +/− 3.2%, 27.5 +/− 3.1%, data are mean +/− SEM; effect of group: *F*_1,72_ = 6.96, *p* = 0.012, effect of time: *F*_2,72_ = 0.15, *p* = 0.86, group-by-time interaction effect: *F*_2,72_ = 0.83, *p* = 0.44; intermediately active cells: week 0 – 4 in WT: 68.1% +/− 4.1; 77.4 +/− 2.8%, 75.6 +/− 2.9%; in APPPS1: 62.2 +/−2.4%, 66 +/− 2.5%, 65 +/− 2.4%; effect of group: *F*_1,72_ = 5.52, *p* = 0.024, effect of time: *F*_2,72_ = 3.47, *p* = 0.036, group-by-time interaction effect: *F*_2,72_ = 0.006, *p* = 0.45; rarely active cells week 0 – 4 in WT: 16 +/− 4.05%, 9.21 +/− 2.4 %, 13.77 +/− 2.7%, in APPPS1: 10.6 +/− 1.8%; 6.3 +/− 1.1 %, 7.5 +/− 1.6%; effect of group: *F*_1,72_ = 3.9, *p* = 0.056, effect of time: *F*_2,72_ = 5.24, *p* = 0.0075, group-by-time interaction effect: *F*_2,72_ = 0.51, *p* = 0.6, WT n = 9, APPPS1 n = 29 experiments, two-way repeated measures ANOVA). **(c-e)** Activity-category specific changes in neuronal activity throughout the 4 week investigation period. **(c)** Highly active neurons in APPPS1 mice underwent a smaller reduction in activity than those neurons in their WT littermates (*p* = 0.046 for week 0-2 (dashed lines); *p* = 0.012 for week 0-4 (solid lines), KS test, WT n = 81 neurons, APPPS1 n = 444 highly active neurons). **(d)** Intermediately active neurons increased their activity on average in APPPS1 compared to WT (*p* < 10^−4^ for week 0-2; *p* < 10^−7^ for week 0-4, KS test, WT n = 312 neurons, APPPS1 n = 984 neurons). **(e)** The activity change of rarely active neurons did not differ between APPPS1 and WT mice (*p* =0.77 week 0-2; *p* = 0.24 week 0-4, KS test, WT n = 94 neurons, APPPS1 n = 177 rarely active neurons). **(f)** Reoccurrence rate over 4 weeks of highly active cells (*p* = 0.025), **(g)** of intermediately active neurons (*p* = 0.019) and **(h)** of rarely active neurons (*p* = 0.3, Mann Whitney Mann U, data are median +/− 95% CI). **(i)** Fraction of intermediately active neurons turning highly active at week 2 (*p* < 10^−5^, Fisher’s exact test). **(j)** Activity gain of novel highly active neurons at week 2 (*p* = 0.1, Mann-Whitney U test, data are median +/− 95% CI). **(k)** Fraction of intermediately active neurons becoming highly active from week 2-4 (*p* < 10^−6^, Fisher’s exact test). **(l)** Activity gain from week 2-4 of those novel highly active neurons (*p* = 0.82, Mann-Whitney U test, data are median +/− 95% confidence interval). KS…Kolmogorov-Smirnov., * *P* < 0.05, ** *P* < 0.01, *** *P* < 0.001

Changes in activity of rarely active neurons, however, did not differ between genotypes (Fig. 3e). As the absolute change in activity is affecting the likelihood to remain within a given category we next compared the stable fraction of neurons within each activity category over time (reoccurrence rate, Fig. 3f-h). In accordance with the change in activity levels, we found that highly active neurons were significantly more likely to remain within the same category in APPPS1 mice (Fig. 3f), while intermediately active neurons were less likely to stay intermediately active over 4 weeks in APPPS1 mice (Fig. 3g). The fraction of consistently rarely active neurons after 4 weeks was close to zero and not different between genotypes (Fig. 3h).

The overall stability of the 3 activity categories throughout the 4 week observation time is based on a largely balanced fractional loss and gain of neurons within each category (see Suppl. Fig. 3). Aberrantly high activity levels, termed hyperactivity of neurons, are considered a hallmark of AD pathology. However, how hyperactivity emerges remains elusive. Does the activity of certain neurons slowly increase over time or is there a sudden pronounced increase in activity detectable in some neurons? To address this question we analysed the population of neurons that only turned highly active at week 2 and 4, respectively (Fig. 3i-l). These ‘novel highly active’ neurons were almost exclusively recruited from former intermediately active neurons in both WT and APPPS1 mice (week 2: WT 79% and APPPS1 94% former intermediate, week 4: WT 96% and APPPS1 97% former intermediately active neurons, Fig. 3a,b). How frequently does this shift in activity occur? While from week 0 – 2 in WT mice 9.7 % of the intermediately active neurons transitioned into highly active, the same was true for 21% of intermediately active cells in APPPS1 mice (Fig. 3i). Those novel highly active neurons at week 2 gained on average 2.48 (1.94 – 3.09 CI) transients/min in WT and 2.03 (1.91 – 2.2 CI) transients/min in APPPS1 mice (Fig. 3j). Interestingly, from week 0 to 2 this effect is not based on a higher average activity within the population of intermediately active cells as both the average activity as well as the activity distribution of intermediately active cells did not differ between WT and APPPS1 mice (median activity intermediately active cells in WT week 0: 2.3 (2.25 – 2.5 CI) transients/min, in APPPS1: 2.48 (2.4 – 2.55 CI) transients/min, *p* = 0.08, Mann Whitney U test; activity distribution KS test *p* = 0.1, data not shown). Similar changes were observed between week 2-4 (Fig. 3k,l) with 7.3% (week 2-4) of intermediately active cells transitioning into highly active neurons in WT and 18.4% (week 2-4) of intermediately active neurons in APPPS1 mice (Fig. 3k). Again, no difference in overall activity gain was detected for those intermediately active cells becoming highly active (Fig. 3l). At the time point week 2, however, the neuronal activity of intermediately active cells was already significantly higher in APPPS1 mice (average neuronal activity intermediately active cells at week 2: WT 2.31 (CI 2.19 – 2.47) transients/min, APPPS1 2.59 (CI 2.49 – 2.69) transients/min, p < 10^−3^ Mann Whitney U test), rendering them more likely to transition into highly active cells.

Thus, “hyperactivity” seems to result from a gradual increase in activity of mainly intermediately active cells over time both in WT and APPPS1 mice and not by a rapid, pronounced transition in activity of individual cells. Importantly, in APPPS1 mice a larger proportion of intermediately active cells transitions into highly active ones.

#### Effect of amyloid plaque proximity

Aβ plaques are a hallmark of AD pathology, and even though their pathogenic relevance is currently debated ^28, 29^, the local plaque environment is associated with a number of pathological features, such as dystrophic neurites, synapse loss and instability ^12^, reactive microglia and astrocytes and hyperactive neurons ^1^. We thus asked whether neurons close to plaques would change their activity more vigorously or frequently than neurons further away from plaques. To this end we measured the distance of each neuron to the nearest plaque, stained by the dye Methoxy-XO 4 in 3D (Fig. 4a,b). In this analysis only experiments, in which we could faithfully track the distance to the nearest plaque for all neurons at all imaging time points were included. Based on the median distance of all neurons to the nearest plaque (Suppl. Fig. 4a, week 0: 39.9μm, week 2: 36.8μm, week 4: 34.9 μm – plaque distance decreases as plaques grow over time ^27, 30, 31^), we divided neurons into close (<= 40 μm form the plaque border) and distant (> 40μm from plaque border) from the nearest plaque, respectively (of note, the effect is also true for other cut offs, such as 20, 60 or 80 μm). The fraction of highly or rarely active neurons did not differ significantly between close and distant neurons at the level of individual experiments (Suppl. Fig. 5a). Relative proportions of highly, intermediately and rarely active neurons in APPPS1 mice were also stable for both close and distant neurons (Fig. 4c). However, neurons close to plaques that were either highly or rarely active were more likely to maintain their activity level than those neurons further away from plaques (Fig. 4d). Intermediately active neurons on the other hand were more stable distant from plaques (Fig. 4d). When analysing the change in neuronal activity for neuronal populations divided into plaque distance bins (neurons located within 0-200m, 20-40mm, 40-60mm, 60-80mm to the closest plaque), we found that the majority of neurons preserve their activity (week 0 – 2: 79%, 82%,78%, 80 %; week 0 – 4: 76 %, 81%, 81%, 86% of the respective bins 0-20, 20-40, 40-60, 60-80um distance change their activity within a range of +/− 2 transients/min), an effect that was independent of the plaque proximity (Fig. 4e). Taken together, our results indicate that neuronal activity at the single cell level varies only slightly, is as stable as in WT mice and is independent from plaque proximity. In addition to overall neuronal activity alterations, pathological changes could also influence the temporal correlation of activity between neurons. We thus investigated the synchrony of the activity of the neurons recorded in a given field of view (Fig. 5a). The Pearson’s correlation coefficient (R) was significantly increased in APPPS1 mice compared to WT mice across all time points, an effect that was not simply based on neuronal activity as the correlation of the shuffled data, was significantly lower (Fig. 5b). We next asked whether functional ensembles with a high level of synchrony are located in spatial vicinity to each other, by testing whether the pairwise correlation was dependent on the physical distance of the neurons (Fig. 5c). Consistent with previous work, we observe a higher correlation between neighbouring neurons (within 20 μm ^32^). In addition, pairwise correlations in APPPS1 at all distances were higher than in WT mice (Fig. 5c). Collectively, our results demonstrate that neurons in the APP transgenic mice exhibit a strongly enhanced level of correlated activity.

**Fig. 4.**
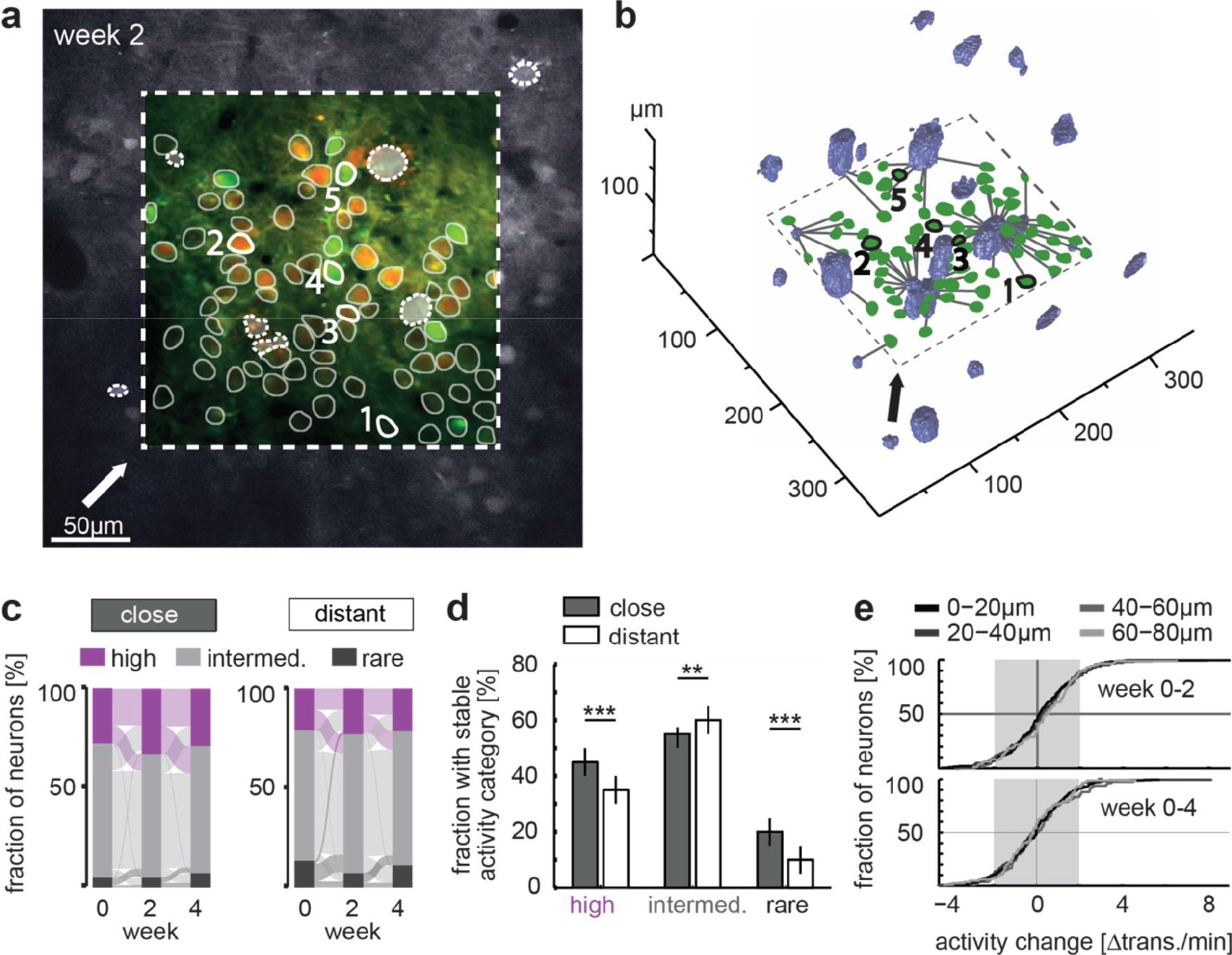
Impact of amyloid plaque proximity on the dynamics of neuronal activity. **(a)** Representative *in vivo* two-photon image, denoting the location of an imaging field of view (dashed rectangle) within the single plane of a z-stack, acquired for the measurements of the distances between plaques and neurons (GCaMP6s green, mRuby2 red). Selected regions of interest (ROIs) and amyloid plaques are marked by solid and dashed lines, respectively. Example ROIs marked by numbers 1-5 correspond to neurons 1-5 in (b). The arrow indicates the viewing angle chosen in (b). **(b)** 3D reconstruction of the imaged area shown in (a) demonstrating the location and proximity of amyloid plaques (blue) and the selected ROIs (green). ROI numbers correspond to the ones in (a). Lines connecting the ROIs and the plaques represent the shortest distance from a given neuron to its nearest plaque border. **(c)** Relative proportions of highly active, intermediately active and rarely active neurons and their fractional change over time in APPPS1 mice for close (<= 40μm from plaque border, left panel) and distant (> 40 μm from plaque border, right panel) neurons (for alluvial plots all neurons were pooled). Relative proportions of the 3 activity categories did not differ significantly and were stable for both close and distant neurons over time (fraction highly active neurons: effect of group: *F*_1,56_ = 0.44, *p* = 0.51, effect of time: *F*_2,56_ = 0.13, *p* = 0.88, group-by-time interaction effect: *F*_2,56_ = 0.78, *p* = 0.46, fraction of intermediately active neurons: effect of group: *F*_1,56_ = 0.18, *p* = 0.68, effect of time: *F*_2,56_ = 0.5, *p* = 0.61, group-by-time interaction effect: *F*_2,56_ = 1.14, *p* = 0.33, fraction rarely active neurons: effect of group: *F*_1,56_ = 0.68, *p* = 0.41, effect of time: *F*_2,56_ = 2.62, *p* = 0.08, group-by-time interaction effect: *F*_2,56_ = 1.16, *p* = 0.32; two-way repeated measures ANOVA, close n = 15, distant n = 15 experiments; 326 neurons close, 326 neurons distant). **(d)** Fraction of neurons in each activity category that remain within the same activity category over the whole imaging period assessed for neurons close (striped) and distant (filled) from plaques (highly active neurons *p* < 10^−4^, intermediately active neurons *p* = 0.003, rarely active neurons *p* < 10^−3^), data are bootstrapped median +/− 95% confidence interval. **(e)** Cumulative distribution of the change in neuronal activity for neurons within different plaque-distance bins. Activity changes are calculated between week 0 - 2 (upper panel, *F*_3,615_ = 0.47, *p* = 0.7, one way ANOVA) and week 0 – 4 (lower panel, *F*_3,615_ = 0.97, *p* = 0.41, one way ANOVA). The vast majority of neurons (> 75%) change their activity within a range of +/− 2 transients/min (gray box). ** *P* < 0.01, *** *P* < 0.001

**Fig. 5.**
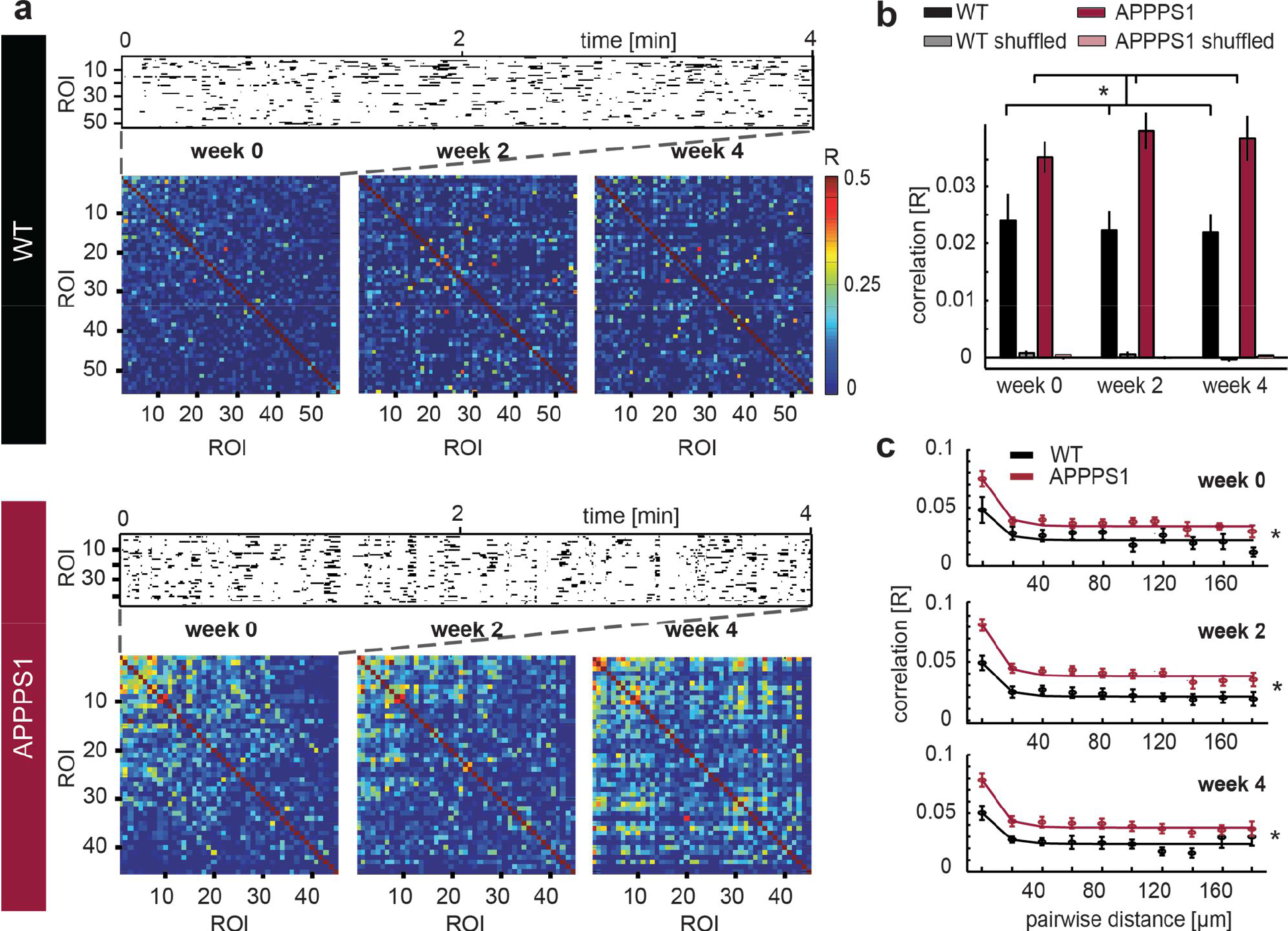
Increased pairwise activity correlation in APPPS1 transgenic mice. **(a)** Representative example of a raster plot depicting binarized neuronal activity (upper panel) in both a WT and an APPPS1 mouse. The pairwise correlations (Pearson’s correlation coefficient R) were computed and sorted at the first time point and displayed as color-coded correlogram for each time point (bottom of each panel) for the WT (upper panel) and the APPPS1 (lower panel) mouse. **(b)** Average neuronal correlation in individual experiments in WT (black) and APPPS1 (red) mice for actual experimental (bright color bars) and shuffled (pale bars) data. The Pearson’s correlation coefficient (R) was significantly increased in APPPS1 mice compared to WT mice (effect of group: *F*_1,72_ = 6.44, *p* = 0.015; effect of time: *F*_2,72_ = 0.97, *p* = 0.38; group-by-time interaction effect: *F*_2,72_ = 0.57, *p* = 0.57) across all time points. The correlation of the shuffled data was significantly lower (WT vs WT shuffled, effect of group: *F*_1,72_ = 43.66, *p* < 10^−9^; effect of time: *F*_2,72_ = 0.18, *p* = 0.83; group-by-time interaction effect: *F*_2,72_ = 0.08, *p* = 0.93, APPPS1 vs APPPS1 shuffled, effect of group: *F*_1,72_ = 167.52, *p* < 10^−9^; effect of time: *F*_2,72_ = 1.41, *p* = 0.25; group-by-time interaction effect: *F*_2,72_ = 1.19, *p* = 0.31, two-way repeated measures ANOVA, WT n = 9 experiments, APPPS1 n = 29, data are mean +/− SEM). **(c)** Neuronal correlation as a function of distance between neuronal pairs in WT (black) and APPPS1 (red) mice for three imaging time points. Solid line represents an exponential fit for visual guidance. At all imaging time points we found higher pairwise correlations in APPPS1 at all neuronal distances (week 0: effect of group *F*_1,2_ = 21.4, *p* = 10^−5^, effect of distance: *F*_2,9_ = 7.42, *p* = 10^−8^, week 2: effect of group *F*_1,239_ = 51.12, *p* = 10^−11^, effect of distance: *F*_9,239_ = 10.17, *p* = 10^−12^, week 4: effect of group *F*_1,239_ = 32.76, *p* = 10^−7^, effect of distance: *F*_9,239_ = 7.41, *p* = 10^−9^, two-way ANOVA, WT n = 7 experiments, APPPS1 n = 23 experiments, only experiments with pairwise distances covering the whole range of distance bins were considered, * *P* < 0.05)

### Discussion

We monitored the activity levels of individual neurons over a time course of 4 weeks in awake APPPS1 transgenic mice and their non-transgenic litter mates. We a) confirm the occurrence of a larger fraction of highly active neurons in awake APPPS1 mice, b) demonstrate that the activity levels in APPPS1 mice vary only slightly over time, to a degree comparable to WT mice, c) particularly intermediately active cells in APPPS1 mice slowly increase their activity over time, which could lead to the development of highly active cells (Suppl. Fig. 5b), d) highly active cells are more likely to stay highly active in APPPS1 and e) altered neuronal activity is associated with a higher pairwise correlation. Our findings thus support the notion of slow progressive network alterations driving aberrant activity patterns, which are likely based on a chronic excitation/inhibition imbalance.

### Aberrant neuronal activity levels are a key feature of AD pathology

Changes in neuronal activity levels, seen both as a strong increase as well as a reduction in firing rates and an intermittent higher degree of neural synchrony are well documented in mouse models and human patients ^1, 2, 16, 22, 33^. Aberrant single cell activity has been shown to impair neural network function ^3, 4^ and is believed to result in cognitive deficits typical of AD. Restoring single cell activity levels and thus neural circuits is thus one of the main therapeutic targets to combat AD. However, the mechanisms driving neuronal and network dysfunction still remain incompletely understood, hampering the development of effective treatment strategies. Earlier work has suggested that Aβ oligomers can directly cause neuronal hyperactivity ^16, 17^, potentially by facilitating glutamate release. Whether these mechanisms are also at play under conditions of chronic exposure to Aβ remains unclear. Neurons are normally endowed with the necessary molecular machinery to maintain their average firing rates upon a perturbation as a homeostatic response ^21, 34^. As such, increased firing rates due to acute exposure to elevated Aβ levels are likely to be a transient phenomenon ^17^. Our data, however, suggests that aberrant activity, in particular high activity levels, are a rather stable single cell feature that is present over many weeks and only emerges slowly. We did observe some fluctuations in the average neuronal activity both in WT and in APPPS1. These fluctuations can even under physiological conditions be as large as an order of magnitude difference in spike rate at a given time scale ^34^ and might very well be brain region and cell-type specific. For this reason the reoccurrence rate of neurons within a given activity category drops over time. In addition to physiological variability, there are methodological constraints largely determined by the recording time. In our experiments we continuously recorded from neurons over ~ 8 min – the detection stringency is thus 0.125 transients/min. To be considered an intermediately active ROI, a neuron would have to fire at least 2 transients during the recording time. As the reoccurrence rate of rarely active cells in both WT and APPPS1 is close to 0 after 4 weeks, we assume that almost all cells we recorded from are in fact active cells. The fraction of rarely active cells was also low and did not significantly differ between WT and APPPS1. On the other hand we did already observe a strong increase in highly active cells during this early phase of amyloid pathology. This finding is in agreement with a previous study, which also provides evidence for activity alterations in cortex as early as 4.5months of age in a similar APP/PS1 transgenic mouse model ^4^. Importantly, most of these earlier studies have been conducted in low-level anesthetized mice upon acute window preparations using synthetic calcium indicators ^1, 4, 16^. Volatile anaesthesia have been shown to strongly reduce particularly the inhibitory drive during sensory processing ^35^ and to trigger network synchrony ^36^. Our data derived from awake mice now further substantiate those earlier studies. As we did not observe an increase in rarely active cells, our data argue for a temporal sequence in which during early stages of the disease neuronal activity is elevated and possibly only at later stages some neurons turn rarely active. In our dataset we at least did not find any compelling evidence for highly active cells becoming rarely active over the 4 week imaging period (number cells highly active at week 0 and rarely active at week 4: WT mice 1 out of 471 and in APPPS1 mice 4 out of 1514 total neurons).

We did, however, find a slight increase in absolute neuronal activity particularly for intermediately active cells, which ties in with a larger fraction of intermediate cells turning highly active. At the same time some of those highly active neurons would decrease their activity levels to be classified as intermediately active (likely based on the expected degree of activity fluctuations (see above)), thereby not yet resulting in a change in the fraction of highly active cells. If the slight increase in average activity (~0.5 transients/min within 2-4 weeks) of intermediately active neurons persisted, it could well serve as a source for future highly active cells. Whether this process is occurring stepwise or gradual with a linear progression needs to be addressed in future studies. Overall this slow process would fit well with progressive network dysfunction based on E/I imbalance underlying the observed alterations in spontaneous activity ^3^. This notion is further corroborated by the observed increased correlation of neuronal activity in APPPS1 mice. Correlated activity argues against cell autonomously elevated activity levels of individual cells but rather point to circuit mechanisms (such as increased synaptic inputs or a lack of inhibition). Interestingly, similar alterations within cortex were also recently described in other mouse models of neurodegenerative diseases (NDs), such as Huntington’s disease ^37^, potentially indicating that impaired E/I balance constitutes a general feature in NDs. Previous work has reported epileptic discharges in APP transgenic mice primarily seen during low gamma states, which the authors linked to impaired function of particularly parvalbumin (PV) positive interneurons ^22^. The increased activity correlation we observed in our awake imaging studies, however, unlikely represents epileptiform discharges as the hypersynchrony underlying these brain activity patterns is expected to result in much higher levels of activity correlation ^38^. Interestingly, recent findings suggest that PV interneurons are also involved in facilitating ‘baseline’ synchrony of neural circuits during quiescent states and that their optogenetic silencing yields a drop in network synchrony both during visual stimulation as well as during spontaneous activity ^39^. The observed increase in correlated spontaneous activity in our data thus might result from aberrant correlated synaptic excitatory inputs, which could result from structural remodelling known to accompany for instance amyloid plaque pathology ^12, 40^. In line with this idea are also findings reporting increased local connectivity and synchronization in AD, while inter-regional connectivity was strongly decreased ^41, 42^. In conclusion, the network mechanisms underlying neural synchronization in AD under different conditions remains elusive thus far and warrants further scrutiny.

### Impact of amyloid plaque proximity

Earlier work has suggested that highly active cells are spatially linked to plaque proximity ^1^. This finding, however, remains controversial as it could not be replicated in other studies ^3, 4^. We thus also investigated neuronal activity levels and their dynamics separately for neurons close to plaques (0-40μm away from the closest plaque border) and those further away. Although there was a trend towards a higher fraction of highly active and a lower fraction of rarely active neurons close to plaques, this effect was not significant at the level of individual experiments (field of views). We did, however, find differences with respect to the stability of the neuronal activity levels. Close to plaques neurons were more likely to remain either highly or rarely active, while distant from plaques more cells would remain intermediately active. These data indicate a ‘solidification’ of aberrant activity levels close to plaques, while neurons further away might still be in a transition phase.

One of the shortcomings of our study is the rather short observation time of 4 weeks. Certainly, a much longer imaging period is desirable. However, due to technical constraints these experiments are currently difficult to realize. Firstly, typically the window quality decays over time due to regrowth or dural thickening. Secondly, the usage of AAV-mediated indicator expression is resulting in nuclear expression of the indicator (so-called ‘filled cells’, ^25^). Filled cells display altered calcium kinetics and response properties ^25^. We can thus not rule out any impact on spontaneous activity levels, for which reason we excluded filled cells from the analyses.

Taken together, our data, to the best of our knowledge for the first time, provide evidence that aberrant neuronal activity is a stable, single cell feature in AD. During the early phase of the disease it develops slowly over time by a gradual increase of the activity of single cells. These data are well in line with the notion of a progressive neural circuit dysfunction, due to an excitation/inhibition imbalance. Future studies should address the dynamics seen in later stages of the disease as well as the changes occurring on dedicated populations of inhibitory interneurons. Another incompletely explored aspect is the fact that homeostatic mechanisms seem to fail in AD. Irrespective of the kind of perturbation, neurons and neural circuits are typically bound to maintain a given average firing rate through processes collectively termed integrated homeostatic network ^43^. Possible reasons underlying an ineffective homeostatic response could involve a failure of the sensing or effecting machinery. Alternatively, an overshooting reaction and thus a maladaptive process could also take place ^44, 45^. These recently emerging concepts warrant further scrutiny as they also pinpoint to downstream circuit alterations that might eventually appear independent of Aβ levels and are thus not amenable to Aβ targeting therapeutic strategies. Our findings therefore also emphasize the need to monitor therapeutic effects by means of longitudinal observations ideally at single cell resolution. Importantly, as aberrantly high activity levels are a stable, single cell feature those cells could on the one hand serve as read-out for the efficacy of novel treatment strategies and moreover, can also be targeted selectively in future therapeutic approaches.

## Materials and Methods

### Animals

All procedures were carried out in accordance with an animal protocol approved by the Ludwig-Maximilians-University Munich and the government of Upper Bavaria (ref number GZ: 55.2-1-54-2532-163-13).

As an AD transgenic model the double transgenic mouse line APPPS1 ^26^ on a C57BL/6J genetic background was used. These mice co-express the mutant amyloid precursor protein (APP, Swedish double-mutation KM670/671NL) and mutant presenilin 1 (PS1, L166P) under the control of the neuron-specific Thy-1 promoter (referred to as APPPS1). The line is hemizygous for both transgenes. As controls in all experiments the non-transgenic (wild type) littermates were used (referred to as WT). Both sexes were used in the study. In total 9 APPPS1 (29 fovs, 1691 neurons) and 5 WT mice (9 fovs, 562 neurons) were included in the study.

Before the cranial window implantation surgery mice were housed in groups of three to six individuals in standard cages, with standard bedding and additional nesting material. After the surgery, they were singly housed in standard cages. Food and water were provided ad libitum. Mice were kept under a 14/10-hr light/dark cycle.

### Surgery

For *in vivo* imaging, a chronic cranial window was implanted as described previously ^46, 47^. Mice underwent surgery at the age of 3.5 months. Briefly, mice were anesthetized by an intraperitoneal injection of Ketamine/Xylazine (14 mg/kg body weight; WDT/Bayer Health Care). Additionally, Dexamethasone (6 mg/kg body weight; Sigma) was intraperitoneally administered immediately before surgery. Firstly, a round craniotomy of 3 mm in diameter was made above the right hemisphere frontal to bregma (coordinates of the center of the craniotomy: 1.5 mm anterior, 1.75mm lateral to bregma) using a dental drill (Schick-Technikmaster C1; Pluraden; Offenbach, Germany). Then the virus injection was performed within the center of the craniotomy. The virus AAV2.1.hSyn1.mRuby2.GSG.P2A.GCaMP6s.WPRE.SV4 (^24^; Cat.No 50942-AAV1 Penn Vector Core) was injected at 1:50 dilution of the original stock (final virus titer 0.33 × 10^13^ GC ml^−1^) at a volume of 300)l each in 3-5 injections at a depth of 0.8 mm, at a speed of 33 nl/min using the NANOLITER 2010 Injector with Micro4 Controller (World Precision Instruments). The injection site was selected such that no big blood vessels would be damaged by the injection. Immediately after all the 3-5 injections were done the cranial window was covered with round coverslip (3mm, 0.16 − 0.19 mm thickness, World Precision Instruments). The coverslip was sealed using dental acrylic (Cyano-Veneer fast; Schein). A custom-made small metal bar was attached with dental cement next to the coverslip to allow for a stable head-fixation during training and awake imaging sessions. After surgery, mice received subcutaneous doses of the analgesic Carprofen (7.5 mg/kg; Pfizer) and the antibiotic Cefotaxime (5 mg/kg; Pharmore).

### Longitudinal awake *in vivo* two-photon imaging

Imaging was performed in awake, head-fixed mice as described previously ^48^. Mice were trained to accommodate to the head-fixation for 14-21 days prior to imaging.

Weekly imaging sessions started four weeks after the surgery to allow mice to recover and accommodate to the setup and the cranial windows to become stable. If the mouse was not getting habituated to the setup and showed signs of distress during fixation after the training period, it was removed from the experimental group.

Around 5-15 hours before each imaging session, Methoxy-X04 (Xcessbio, San Diego, CA, USA, 3.3% vol of 10 mg/ml stock solution in DMSO (light shielded), 6.66% vol Cremophore EL (Sigma Aldrich) in 90% vol PBS), was intraperitoneally injected at a concentration of 3.33 mg/kg body weight to stain amyloid plaques *in vivo* ^49^. Before each imaging session, mice were head-fixed and placed under the microscope for 5 min to habituate. Imaging was performed in the dark without any additional stimuli. In each mouse, two to six regions at depths of 120 – 200 μm below the pial surface (layer 2/3) of the frontal cortex were imaged. *In vivo* time-lapse imaging stacks were acquired at a frame rate of 10 Hz using the LaVision Trim Scope microscope equipped with 2 tunable Ti:sapphire two-photon lasers (Coherent Chameleon and Mai Tai Spectra Physics). The setup was controlled using LaVision Imspector software (LaVision Biotech, Germany). The Chameleon laser was tuned to 940nm, which enabled simultaneous excitation of mRuby2 and GCaMP6s. A 25×, NA 1.05 water-immersion objective (Olympus) was used. For the first cohort the imaging dimensions were 173 × 173 pixels; corresponding to 151 μm × 151 μm, for the second cohort − 223 × 223 pixels and 220 μm × 220 μm. At each session the same cells for each area of interest were imaged over 5000 frames (8.3 min duration).

For the analysis of plaque distances z-stacks capturing the surrounding tissue of the imaged area were acquired, with each stack covering 250-350 μm in depth (520 × 520 pixels; x,y dimensions: 350 μm, z increments 0.5μm). The simultaneous excitation of Methoxy-X04 was achieved by the Mai Tai laser tuned to 750 nm. During the z-stack acquisition mice were anesthetized by 0.5 vol % isoflurane subsequent to the awake imaging session. At all times laser power was kept below 80mW measured at the back-focal plane of the objective. Emitted fluorescence light was split at 495 nm and 560nm, to separate the emitted light from Methoxy-XO4, GCaMP6s and mRuby2 and detected by photomultiplier tubes. During head-fixation, mice typically showed long episodes of quiet wakefulness (quiet phase) interrupted by brief episodes of whisking or grooming (active phase). We recorded the behaviour of the mouse by a webcam, controlled by the LaVision Imspector software, which allowed for a synchronisation of the two-photon and the behavioural recordings. Whisking/grooming epochs were not considered in the analysis of neuronal activity.

### Immunohistochemical identification of inhibitory neurons

The PFA-fixed brains were cut on a vibratome (Leica VT 1000S) into 50 μm thick coronal sections. Immunohistochemistry was performed on free-floating sections

Sections were incubated overnight in 2% Triton X-100 in PBS at room temperature for permeabilization of the tissue, then blocked for 2 hours at room temperature with 3% I-Block™ Protein-Based Blocking Reagent (Thermo Fisher Scientific) containing 0,2% Triton X-100 in PBS, before they were incubated with the primary antibodies (mouse anti-GAD67 in 1:500 dilution (Millipore, catalogue number MAB5406B)) overnight. Incubation with the secondary antibody (goat anti-mouse Alexa Fluor 647 1:500, Life Technologies, catalogue number A21236) was conducted for 2 days at room temperature. Confocal stacks (Zeiss LSM 780) were acquired first at a low magnification (10x objective) to allow for identification of virus-transfected regions. Confocal stacks of re-identified imaging spots were then acquired with a 40x objective. Analysis of images was performed using ZEN software (Zeiss) in raw z-stacks by manually scrolling through respective frames and marking mRuby2- and GAD67-positive neurons.

### Image data processing and analysis

Collected images were processed and analysed using custom written routines in MATLAB (The MathWorks, Inc.) and ImageJ (http://rsb.info.nih.gov/ij/). Image pre-processing for calcium imaging data was done by custom-made script written by P. Goltstein ^32^. In brief, in vivo two-photon recordings were corrected for slight brain displacement artefacts in the x-y plane by realigning the images ^50^. Recordings with clear movement artefacts along the z-axis were excluded from the further analysis. Regions of interest (ROIs) were outlined semi-automatically based on a maximum projection of all frames for all neurons with the help of custom-made GUI for each repetition block separately (see https://github.com/JorritMontijn/Preprocessing_Toolbox). Indicator-filled cells – seen as bright cells in the GCaMP6 channel without the nucleus being spared at all times of the recordings – were excluded from analyses. The fluorescent intensity of all pixels contained within a given ROI was averaged for each frame for both channels. To correct for contamination by the neuropil, a region surrounding the selected ROI was selected and the average fluorescent intensity within that area was calculated for each frame. The corrected ROI signal was computed based on the equation ^3, 25^:

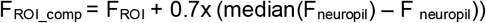

F_ROI_comp_ represents the actual signal within the selected ROI after compensating for neuropil contamination, while F_ROI_ reflects the signal within the initially selected ROI. F_neuropil_ corresponds to the signal within the surrounding neuropil. Traces were next low pass filtered at 5Hz and slow fluctuations removed by subtracting the 8th percentile within a window of +/−15 second ^51^. In order to estimate F0 we subtracted the 8th percentile in a very short window of +/−15 seconds and used the median of all values below the 60th percentile of this ‘noise band’ as F0. This procedure allowed for reasonable F0 detection for both highly and rarely active cells. To classify a neuron as active, it had to display at least one prominent transient exceeding the threshold of F0 plus 3× the standard deviation of the noise band for more than 9 frames (equaling 1 second). Transients were identified on traces smoothed over 5 frames and had a minimum distance of 15 frames (1.5 seconds) and a minimum height of 3 × standard deviation of the noise band. The classification into activity groups is based on the frequency of transients: rarely active 0- < 0.25 transients/min; intermediately active 0.25 − 4 transients/min; highly active > 4 transients/min ^1^. The correlation (Pearson’s R) of the activity between ROIs was based on binary traces. To this end the traces were smoothed across 20 frames (2 seconds) and binarized at a threshold of 2× standard deviation. The distance between cells reflects the distance of the ROIs’ centroids.

### Plaque distance analysis

The analysis of the amyloid plaque distances was carried out in MATLAB (MathWorks) using custom written routines. The position and size of the ROIs were projected into the 3D rendered overview stack. The two channels carrying either the mRuby2 or the Methoxy-XO4 signal were background subtracted, and subsequently the GCaMP6s channel was subtracted from the Methoxy-XO4 channel in order to remove slight bleed through. Each frame of the Methoxy channel was median filtered and binarized using the background plus 3× standard deviation as threshold. Plaque distance is the 3D Euclidean distance between the centroid of the respective ROI and the nearest Methoxy positive voxel. All overview stacks were visually checked for correct plaque detection and 3D rendered stacks were inspected for accurate neuron - plaque assignment (ruling out accidental pairing with voxels carrying signal stemming from the dura (due to generation of second harmonics)). The distances resulting from an incorrect assignment of plaque voxels were exchanged by manually measured distances. Manual measurement was done between the centroid of the respective ROI and the nearest Methoxy positive voxel using the manual measurement tool in ImageJ (http://rsb.info.nih.gov/ij/), taking into account the Pythagorean Theorem in case the nearest plaque was not in the same plane as the ROI.

### Statistics

If not stated otherwise in the text, neuronal activity and fractions of different activity categories and their changes over time were compared using a two-way repeated measure ANOVA. Distributions of activity changes were compared using a Kolmogorov-Smirnov (KS) test. P-values are reported as follows: *P < 0.05, **P < 0.01 and ***P < 0.001.

## Acknowledgements

This work was funded by the Deutsche Forschungsgemeinschaft (DFG, German Research Foundation) under Germany’s Excellence Strategy within the framework of the Munich Cluster for Systems Neurology - EXC 2145 SyNergy – ID 390857198 (VK, JH, SL), the DFG, Emmy Noether Programme (SL), the Deutsche Gesellschaft für Muskelkranke (SL) and the Graduate School for Systemic Neurosciences GSN-LMU (VK). We are thankful to Sonja Blumenstock for providing the in vivo setup scheme in Figure 1A and Finn Peters for support in MatLab programming. We also thank Eric Grießinger, Sarah Hanselka, and Nadine Lachner for excellent technical support. We are grateful to the Genetically-Encoded Neuronal Indicator and Effector (GENIE) Project and the Janelia Farm Research Campus of the Howard Hughes Medical Institute, in particular to Vivek Jayaraman, Rex A. Kerr, Douglas S. Kim, Loren L. Looger, and Karel Svoboda for developing and distributing the genetically encoded calcium indicator GCaMP6s. We are also grateful to Mathias Jucker for providing the APPPS1 mice.

## Author Contributions

Conception and design of study (VK, PM, SL, JH), data acquisition (VK, PM), software development (PG, SL), data analysis (VK, SL), data interpretation (VK, SL), manuscript preparation (VK, SL, help from all authors), securing funding (JH) and project supervision (JH, SL).

## Supplementary Material

**Suppl. Fig. 1.**
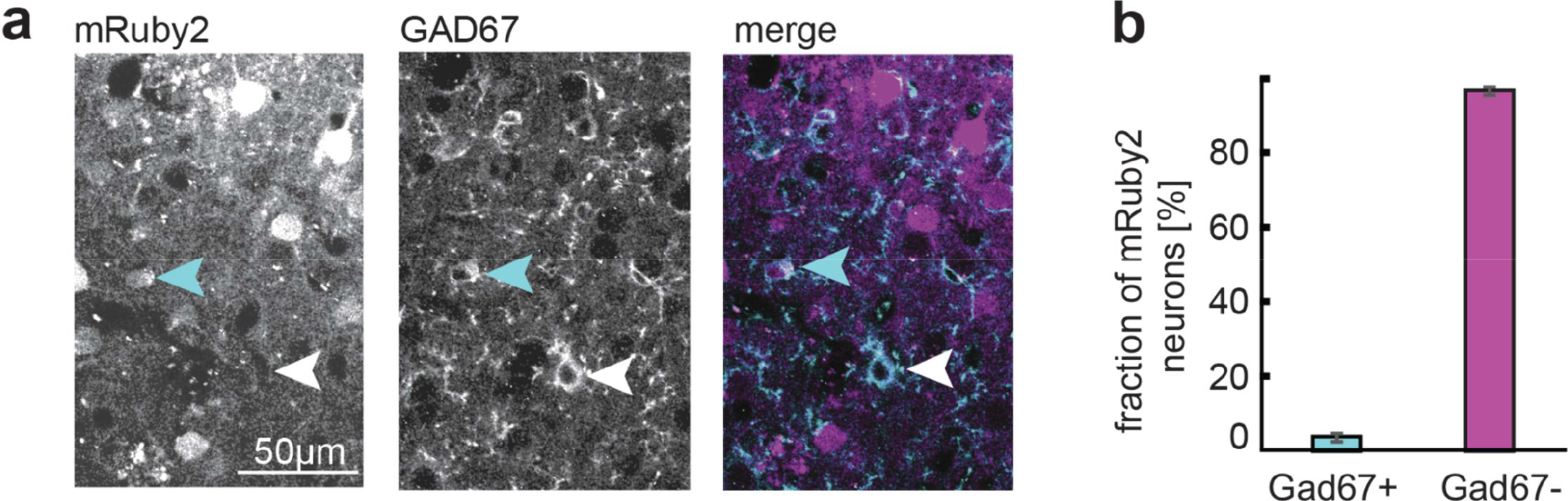
Recorded neurons are mainly excitatory. **(a)** Immunohistochemical analysis of GCaMP6s-mRuby2-expressing neurons. Left, middle and right images represent confocal images of mRuby2, Gad67 immunohistochemical staining and merged image, respectively. White arrowhead marks a Gad67 positive neuron, cyan arrowhead points to an mRuby2 positive, Gad67 positive neuron. **(b)** Quantification of the relative proportion of Gad67 positive, mRuby2 expressing neurons. 3.57% of the mRuby2 expressing neurons (which co-express GCaMP6s) were positive for Gad67 (760 mRuby2-positive neurons, 3 mice, data are mean +/− SD)

**Suppl. Fig. 2.**
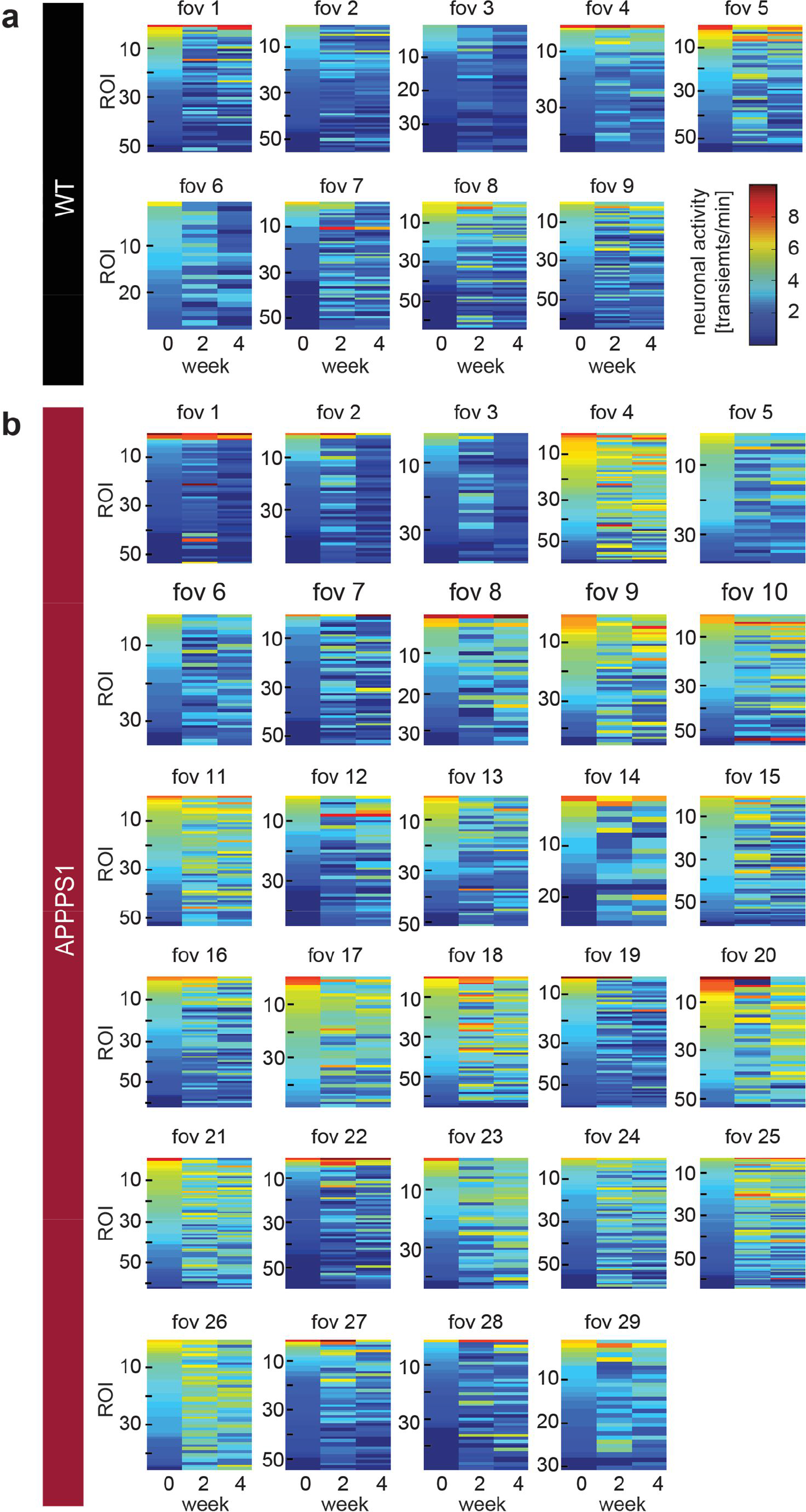
Activity of individual neurons over time. **(a-b)**. Heat maps depicting color-coded activity of individual ROIs (each row represents a single ROI) over time, sorted at the first time point and the order is maintained for all imaging time points for each FOV in WT (a) and APPPS1 mice (b)

**Suppl. Fig.3.**
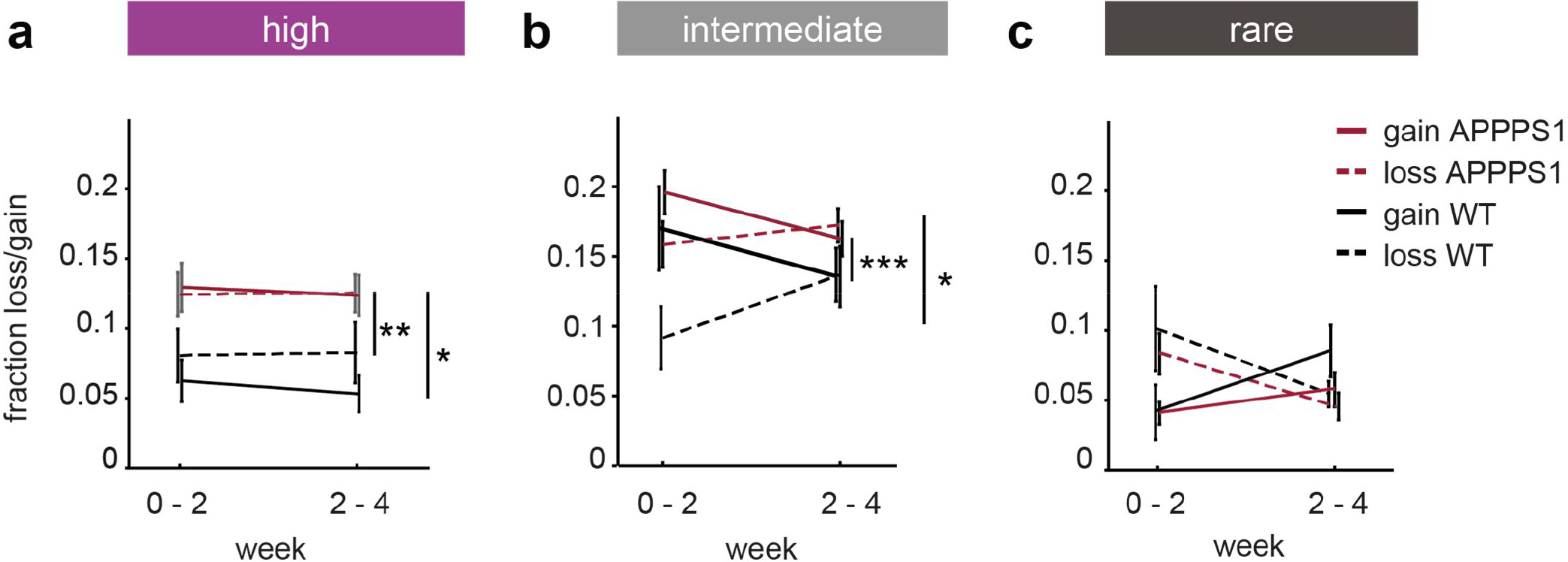
Loss and gain within activity categories. **(a)** Fraction of gained (solid lines) and lost (dashed lines) highly active neurons in WT (black) and APPPS1 (red) mice (fraction lost highly active cells WT vs APPPS1: effect of group F_1,38_ = 4.65, *p* = 0.037, effect of time F_1,38_ = 0, *p* = 0.95, group-by-time interaction effect F_1,38_ = 0, *p* = 0.97; fraction gained highly active cells WT vs APPPS1: effect of group F_1,38_ = 9.77, *p* = 0.0034, effect of time F_1,38_ = 0.22, *p* = 0.64, group-by-time interaction effect F_1,38_ = 0.01, *p* = 0.91; gain vs loss in WT: effect of group F_1,20_ = 1.76, *p* = 0.19, effect of time F_1,20_ = 0.04, *p* = 0.83, group-by-time interaction effect F_1,20_ = 0.11, *p* = 0.74; gain vs loss in APPPS1: effect of group F_1,56_ = 0.01, *p* = 0.92, effect of time F_1,56_ = 0.04, *p* = 0.85, group-by-time interaction effect F_1,56_ = 0.06, *p* = 0.81). **(b)** Fraction of gained (solid lines) and lost (dashed lines) intermediately active neurons in WT and APPPS1 mice (fraction lost intermediately active cells WT vs APPPS1: effect of group F_1,38_ = 6.37, *p* = 0.016, effect of time F_1,38_ = 2.1, *p* = 0.16, group-by-time interaction effect F_1,38_ = 0.83, *p* = 0.37; fraction gained intermediately active cells WT vs APPPS1: effect of group F_1,38_ = 15.44, *p* = 0.0003, effect of time F_1,38_ = 9.45, *p* = 0.0039, group-by-time interaction effect F_1,38_ = 4.13, *p* = 0.05; gain vs loss in WT: effect of group F_1,20_ = 2.08, *p* = 0.16, effect of time F_1,20_ = 0.07, *p* = 0.79, group-by-time interaction effect F_1,20_ = 3.93, *p* = 0.06; gain vs loss in APPPS1: effect of group F_1,56_ = 1.1, *p* = 0.3, effect of time F_1,56_ = 0.43, *p* = 0.51, group-by-time interaction effect F_1,56_ = 2.45, *p* = 0.12). **(c)** Fraction of gained (solid lines) and lost (dashed lines) rarely active neurons in WT and APPPS1 mice (fraction lost rarely cells WT vs APPPS1: effect of group F_1,38_ = 0.51, *p* = 0.48, effect of time F_1,38_ = 8.84, *p* = 0.005, group-by-time interaction effect F_1,38_ = 0.1, *p* = 0.76; fraction gained rarely active cells WT vs APPPS1: effect of group F_1,38_ = 0.69, *p* = 0.41, effect of time F_1,38_ = 5.77, *p* = 0.021, group-by-time interaction effect F_1,38_ = 1.45, *p* = 0.24; gain vs loss in WT: effect of group F_1,20_ = 0.42, *p* = 0.52, effect of time F_1,20_ = 0.01, *p* = 0.94, group-by-time interaction effect F_1,20_ = 5.95, *p* = 0.024; gain vs loss in APPPS1: effect of group F_1,56_ = 1.35, *p* = 0.25, effect of time F_1,56_ = 1.26, *p* = 0.27, group-by-time interaction effect F_1,56_ = 8.59, *p* = 0.005, two-way repeated measures ANOVA). Data are mean +/−SEM; * *p* < 0.05, ** *p* < 0.01, *** *p* < 0.001

**Suppl. Fig. 4.**
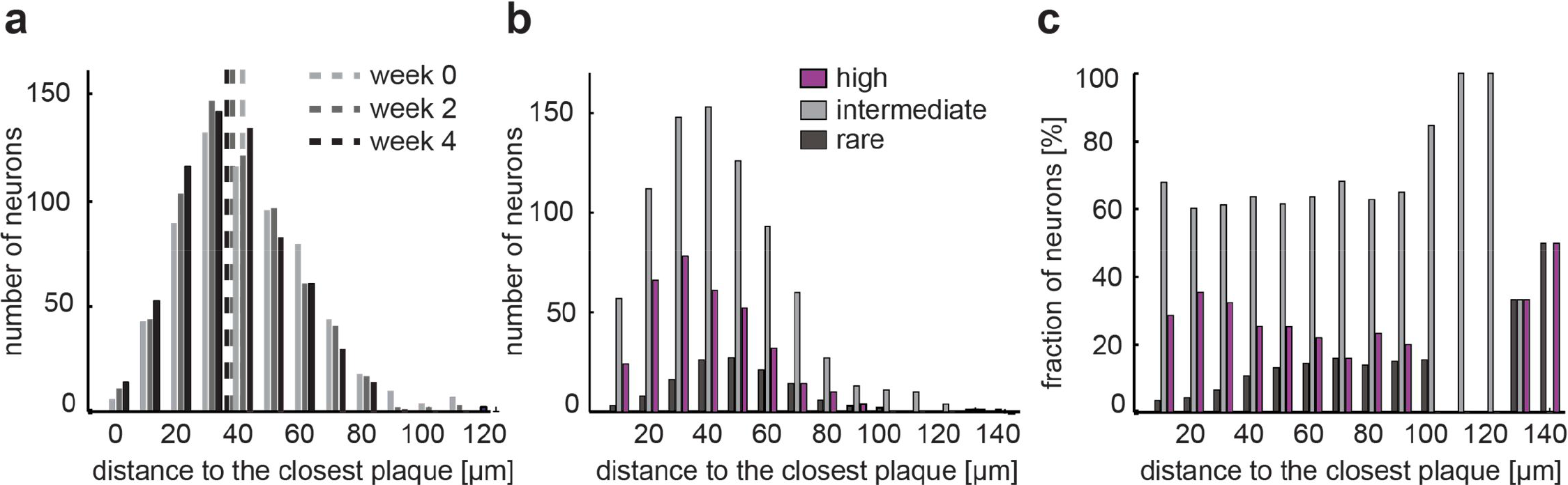
Distribution of plaque distances. **(a)** Distribution of the distances between neurons and their closest plaque at the three imaging time points (median shown as dashed line). **(b)** Distribution of the absolute number of neurons within the three neuronal activity categories at the first imaging time point as a function of plaque distance. **(c)** Distribution of the relative fraction of neurons for the three activity categories at the first imaging time point as a function of plaque distance (data is based on pooled neurons from 15 experiments). Of note, beyond 100 μm very few cells were found, which affects the proportional abundance of the three categories

**Suppl. Fig. 5.**
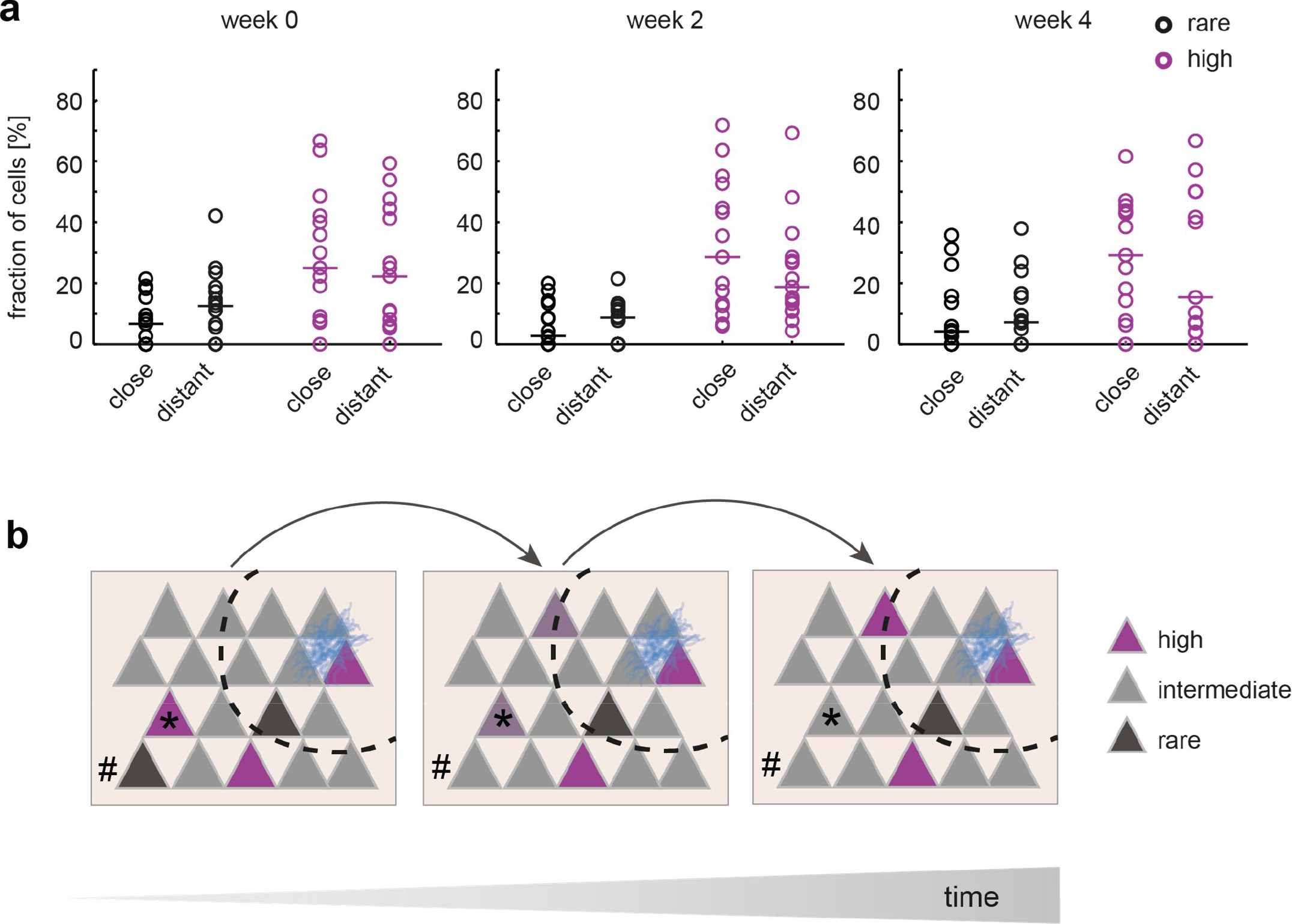
Fractions of rarely and highly active neurons close and distant from amyloid plaques. **(a)** The fraction of highly and rarely active neurons did not differ between close and distant neurons at the level of individual experiments (week 0: fraction rarely active close vs. distant *p* = 0.25, fraction highly active close vs. distant *p* = 0.63; week 2: fraction rarely active close vs. distant *p* = 0.78, highly active close vs. distant *p* = 0.38; week 4: fraction rarely active neurons close vs. distant *p* = 0.63, fraction highly active close vs. distant *p* = 0.93; horizontal line denotes the median. close n = 15 experiments, distant n = 15 experiments, Mann-Whitney U test). **(b)** Scheme illustrating key aspects in the evolution of aberrant neuronal activity during the early phase of AD pathology. Neuronal activity levels in cortex are already altered during the early phase of amyloid plaque deposition (amyloid plaque depicted in blue), resulting in a higher fraction of highly active neurons (magenta). Activity levels remain largely constant over extended periods of time. A proportion of intermediately active neurons (gray) slowly increase their activity thereby turning into highly active neurons over time (example marked by arrow). Distant from plaques (dashed line) aberrant activity (both rarely (#) and highly (*) active cells) is less likely to persist compared to the immediate plaque vicinity

